# IQ-TREE 2: New models and efficient methods for phylogenetic inference in the genomic era

**DOI:** 10.1101/849372

**Authors:** Bui Quang Minh, Heiko Schmidt, Olga Chernomor, Dominik Schrempf, Michael Woodhams, Arndt von Haeseler, Robert Lanfear

**Author notes:** Co-senior authors (Arndt von Haeseler, Robert Lanfear).

## Abstract

IQ-TREE (http://www.iqtree.org) is a user-friendly and widely used software package for phylogenetic inference using maximum likelihood. Since the release of version 1 in 2014, we have continuously expanded IQ-TREE to integrate a plethora of new models of sequence evolution and efficient computational approaches of phylogenetic inference to deal with genomic data. Here, we describe notable features of IQ-TREE version 2 and highlight the key advantages over other software.

---

IQ-TREE is a widely used and open-source software package for phylogenetic inference using the maximum likelihood (ML) criterion. The high performance of IQ-TREE results from the efficient integration of novel phylogenetic methods that improve the three key steps in phylogenetic analysis: fast model selection via ModelFinder (Kalyaanamoorthy et al. 2017), an effective tree search algorithm (Nguyen et al. 2015), and a novel ultrafast bootstrap approximation (Minh et al. 2013; Hoang et al. 2018). Zhou et al. (2018) independently showed that the tree search algorithm in IQ-TREE exhibits good performance in terms of both computing times and likelihood maximization when compared to other popular ML phylogenetics software such as RAxML (Stamatakis 2014) and PhyML (Guindon et al. 2010). IQ-TREE also plays a vital role in the software ecosystem for biomedical research. For instance, it is an integral component of many popular open-source applications such as Galaxy (Afgan et al. 2018), Nextstrain (Hadfield et al. 2018), OrthoFinder (Emms and Kelly 2015), and QIIME 2 (Bolyen et al. 2019).

Since the release of IQ-TREE version 1.0 in 2014, we have continuously developed IQ-TREE to integrate a plethora of new evolutionary models and efficient methods for analyzing large phylogenomic datasets. Here we present IQ-TREE version 2, and highlight the key new features and improvements. To demonstrate its performance and compare it to other software, we analyse two large sequence alignments: a DNA alignment of 110 vertebrate species and 25,919 sites (Fong et al. 2012) which we call the DNA-dataset, and an amino acid alignment of 76 metazoan species and 49,388 sites (Whelan et al. 2017) which we call the AA-dataset.

## Time-reversible models of sequence evolution

IQ-TREE 2 provides a very large collection of more than 200 time-reversible evolutionary models and it supports all standard substitution models for DNA, protein, codon, binary, and multistate-morphological data (Felsenstein 2004; Lemey et al. 2009). Rate heterogeneity across sites can be accommodated either by the discrete Γ distribution (Yang 1994b) with invariant sites (Gu et al. 1995) or a distribution-free rate model (Kalyaanamoorthy et al. 2017). For single nucleotide polymorphism or morphological data the absence of invariant sites can be accounted for by an ascertainment bias correction (Lewis 2001).

IQ-TREE 2 offers a number of advanced models for phylogenomic data including partitioned models (Lanfear et al. 2012; Chernomor et al. 2016), mixture models (Le et al. 2008; Le and Gascuel 2010; Le et al. 2012), posterior-mean site frequency models (Wang et al. 2017), and heterotachy models (Crotty et al. 2019). For allele frequency data IQ-TREE 2 implements the polymorphism-aware models (Schrempf et al. 2016; Schrempf et al. 2019). With partitioned models one can specify either a partition file (as in IQ-TREE 1) or a directory of single-locus alignments. In the latter case IQ-TREE 2 will load and concatenate all alignments within the directory, eliminating the need for users to manually perform this step. The mixture model implementation in IQ-TREE 2 goes beyond that in the PhyML-mixtures (Le et al. 2008) and RAxML-NG software (Kozlov et al. 2019) because it allows for user-defined mixture models. IQ-TREE 2 is also substantially faster than PhyML-mixtures: optimisation of the model parameters for the LG4X model on the AA-dataset took 1.8 minutes in IQ-TREE 2, 1.9 minutes in RAxML-NG, and 67.6 minutes in PhyML-mixtures.

Because substitution models assume time-reversibility, IQ-TREE 1 only infers *unrooted* trees (Felsenstein 1981). In IQ-TREE 2 we included non-time-reversible models, meaning that IQ-TREE 2 can infer *rooted* trees.

## Non-reversible substitution models

IQ-TREE 2 allows users to reconstruct rooted trees using non-reversible models, a feature not available in most ML packages due to numerical and computational expense. We substantially revised the IQ-TREE code to overcome these obstacles. First, due to non-reversibility of the rate matrix *Q*, the naïve computation of the transition probability matrix *P*(*t*) = *e*^*Qt*^, where *t* is the branch length, is unstable due to complex eigenvalues of *Q* (Moler and Loan 1978). Eigen-decomposition and scaling-squaring techniques, e.g., as provided in the Eigen3 library (Guennebaud et al. 2010) remove the numerical problems. IQ-TREE 2 employs eigen-decomposition to diagonalize *Q* into its (complex) eigenvalues, eigenvectors and inverse eigenvectors, which are used to compute *P*(*t*). If *Q* is not diagonalizable, then IQ-TREE 2 switches to the scaling-squaring technique to compute *P*(*t*). The eigen-decomposition is fast but sometimes unstable, whereas the scaling-squaring is slow but stable. Second, IQ-TREE 2 uses a rooted tree data structure and an adjusted pruning algorithm for computing likelihoods of non-reversible models on rooted trees (Boussau and Gouy 2006). Third, we adapted the hill climbing nearest neighbour interchange (NNI), part of the tree search heuristic in IQ-TREE to account for rooted trees. Fourth, we introduced a root search operation that moves the root to the neighbouring branches (by default up to 2 branches away from the current root branch) and retains the rooting position with the highest likelihood. Users can increase this parameter (--root-dist option) to test for the position of the root across more or all branches.

Based on these improvements, we could efficiently implement 99 non-reversible DNA models known as Lie Markov models (Woodhams et al. 2015), the unrestricted model (UNREST) for DNA (Yang 1994a) and, for the first time, the general non-reversible model for amino acid sequences that we call NONREV. For the DNA-dataset (Fong et al. 2012) tree search under the general time-reversible (GTR) model took 2.8 minutes using 4 CPU cores, whereas the UNREST model took 10.7 minutes. For the AA dataset (Whelan et al. 2017) tree search under the protein GTR model took 5.5 hours, whereas the NONREV model took 14.8 hours. The implementation of non-reversible models in IQ-TREE 2 opens new avenues of evolutionary research.

## Fast likelihood mapping analysis

IQ-TREE 2 provides a fast and parallel implementation of the likelihood (quartet) mapping (Strimmer and von Haeseler 1997) to visualize phylogenetic information in alignments or to study the relationships of taxon-groups in large datasets. To this end IQ-TREE 2 evaluates the exact ML value of all relevant quartets. When doing the same with the original implementation of likelihood mapping in TREE-PUZZLE (Schmidt et al. 2002) (i.e. with 10,000 random quartets and exact ML quartet evaluation) for the DNA-dataset it took 282 minutes, whereas the improved implementation in IQ-TREE 2 required only 1 minute using one CPU core and 21 seconds using four cores. Similarly, analysis of the AA-dataset took 60 hours in TREE-PUZZLE, 16 minutes in IQ-TREE 2 with one CPU core, and 4 minutes using four cores. Together with the extended repertoire of sequence evolution models, likelihood mapping facilitates a thorough investigation of much larger sequence alignments.

## New options for tree search

IQ-TREE 2 allows users to perform a constrained tree search (-g option), such that the resulting ML tree will respect a set of user-defined splits, which may also contain polytomies. This option is helpful to enforce the monophyly of certain groups. However, users are advised to always perform the normal unconstrained tree search and then a tree test (see below) to ensure that the constrained tree is not significantly worse than the unconstrained tree.

IQ-TREE 2 also provides a fast tree search (-fast option) that resembles the FastTree2 algorithm (Price et al. 2010). Here, IQ-TREE 2 will compute two starting trees using Maximum Parsimony and Neighbor-Joining (Gascuel 1997), which are then optimized by the hill climbing NNI moves. For our example DNA dataset, the default tree search took 27 minutes, whereas the fast IQ-TREE search and FastTree2 needed 82 and 85 seconds, respectively. For the AA dataset, the default tree search took 5.8 hours, whereas the fast IQ-TREE search and the FastTree2 search took 13.7 minutes and 2.6 minutes respectively. We note that FastTree2 was shown to produce substantially worse trees than RAxML, PhyML and IQ-TREE (Zhou et al. 2018). Hence, this fast option is only recommended for a rough overview of the tree or when the tree topology is not critical for the analysis.

IQ-TREE 2 also provides a --runs option, that conducts multiple independent tree searches, summarizes the resulting trees, and reports the tree(s) with the highest log-likelihood. This option is recommended for difficult datasets, e.g., with many taxa and/or limited phylogenetic signal, to increase the probability that the true ML tree is found.

## Systematically accounting for missing data

Missing data, when some loci or sites are absent for some species, are almost unavoidable in phylogenomic datasets. Missing data can create phylogenetic terraces (Sanderson et al. 2011), where two or more species trees induce the same set of single-locus trees, leading to identical likelihoods under an edge-unlinked partitioned model. IQ-TREE 2 employs the terraphast library (Biczok et al. 2018) to automatically report if the inferred ML tree resides on a terrace. If this is the case, users are advised to reduce the amount of missing data, e.g., by sequencing more loci or filtering out gappy taxa/loci. Moreover, IQ-TREE 2 generalizes the terrace concept to partial terraces (Chernomor et al. 2015), where species trees have only a subset of identical locus trees. IQ-TREE 2 exploits partial terraces to improve tree search under partitioned models and achieves up to 4.5 and 8-fold speedups compared with IQ-TREE 1 and RAxML, respectively (Chernomor et al. 2016).

## Single-locus tree inference

IQ-TREE 2 allows users to infer individual locus trees (-S option), which can be used for subsequent coalescent (e.g., Mirarab et al. 2014) or concordance analyses (e.g., Minh et al. 2018). To this end, users need to either specify a partition file that delineates the borders between loci, or a directory containing the individual locus alignments in individual files. In both cases IQ-TREE 2 will perform separate model selection and tree searches for each locus, which are automatically scheduled on the different CPU cores. This parallelization speeds up computations considerably.

## Fast branch tests

IQ-TREE 2 provides fast and parallel implementations for several existing branch tests including the approximate likelihood ratio test (aLRT) (Anisimova and Gascuel 2006), the Shimodaira-Hasegawa-like aLRT (SH-aLRT) (Guindon et al. 2010), and the aBayes test (Anisimova et al. 2011). The SH-aLRT is parallelized over the bootstrap samples to maximize load balance and efficiency. These tests can be performed on the reconstructed ML tree or a user-defined tree. For the DNA-dataset, PhyML version 3.3.20190321 (Guindon et al. 2010) took 33.9 hours to perform the SH-aLRT, whereas IQ-TREE 2 needed only 1.5 minutes (1,300-fold speedup). On the AA-dataset, PhyML took 173 hours and 27 minutes to perform the SH-aLRT tests, while IQ-TREE 2 needed just 4.5 minutes (a greater than 2000-fold speedup).

## Fast topology tests

IQ-TREE 2 also provides fast and parallel implementations of existing tree topology tests including the Shimodaira-Hasegawa test (Shimodaira and Hasegawa 1999), the approximately unbiased (AU) test (Shimodaira 2002), and the expected likelihood weight (ELW) (Strimmer and Rambaut 2002). Moreover, IQ-TREE 2 is well suited for partitioned models because it provides the site, gene and gene-site bootstrap resampling schemes (Hoang et al. 2018).

For the DNA-dataset CONSEL (Shimodaira and Hasegawa 2001), the original and un-parallelized implementation of the AU test, took 5 minutes to test 100 tree topologies, whereas IQ-TREE 2 took 1.8 minutes with one CPU core and 1.1 minutes with four CPU cores. For the AA-dataset, CONSEL took 9.4 minutes and IQ-TREE 2 took 8.7 minutes with one CPU core and 2.4 minutes with four CPU cores. IQ-TREE offers added convenience because it calculates site log-likelihoods itself, rather than relying on site log-likelihood output provided by other software.

## Scalability with large datasets

IQ-TREE 2 implements several features to facilitate the analysis of large datasets. It uses multi-threading to speed up computations in a range of areas, and can automatically determine the best number of threads for the computer at hand. IQ-TREE 2 periodically writes a compressed checkpoint file that allows to resume an interrupted analysis. IQ-TREE 2 also offers a memory saving mode (-mem option) (Izquierdo-Carrasco et al. 2012), that is automatically invoked when the memory requirement exceeds the RAM size, and a safe mode (-safe option) to avoid numerical underflow for taxon-rich alignments (automatically invoked for datasets with >2000 sequences).

## Documentation, user support and workshop materials

An extensive user manual, quick start guide, tutorials, and command reference are available (http://www.iqtree.org/doc). We actively maintain a forum (https://groups.google.com/d/forum/iqtree) for user support, bug reports and feature requests. We regularly teach IQ-TREE at the Workshop on Molecular Evolution (http://molevol.mbl.edu/), the Workshop on Virus Evolution and Molecular Epidemiology (https://rega.kuleuven.be/cev/veme-workshop/) and the Workshop on Phylogenomics (http://evomics.org). Most workshop materials are freely available at http://www.iqtree.org/workshop/.

## Acknowledgements

The authors wish to thank Suha Naser-Khdour, Cassius Manuel Perez de los Cobos Hermosa, and Lam-Tung Nguyen for fruitful discussions, Lukasz Reszczynski for integrating the terraphast library, Ian Brennan for the logo design, and more than 100 users for helpful feedback and bug reports. Their names are listed in the full release notes (http://www.iqtree.org/release). This work was supported by the Austrian Science Fund [grant no. I-2805-B29] to A.v.H and O.C. and the ANU Futures grant for R.L.

