## Supplement for "IQ-TREE 2: New models and efficient methods for phylogenetic inference in the genomic era"

### Analysis workflow for IQ-TREE 2 paper

This document shows the command lines to run IQ-TREE 2 and other software for the IQ-TREE 2 manuscript for reproducibility purpose.

#### Non-reversible models

This section benchmarks the general time-reversible and non-reversible models implemented in IQ-TREE 2.

For DNA analyses:

```
iqtree -s alignment.phy -T 4 --prefix GTR -m GTR
iqtree -s alignment.phy -T 4 --prefix UNREST -m UNREST
```

For Amino acid analyses:

```
iqtree -s alignment.phy -T 4 --prefix GTR20 -m GTR20
iqtree -s alignment.phy -T 4 --prefix NONREV -m NONREV
```

#### Mixture models

This section benchmarks the LG4X model implementation in PhyML-mixtures, RAxML-NG versus IQ-TREE 2. This only applies to AA analyses.

Run the IQ-TREE 2 analysis:

```
iqtree -s alignment.phy -T 4 --prefix mixture -m LG4X    -te GTR20.treefile
```

Run the PhyML analysis:

```
Phyml-4X_linux64 -i alignment.phy -u GTR20.treefile -o lr -d aa -m LG4X
```

We downloaded the RAxML-NG version 0.9.0 (https://github.com/amkozlov/raxml-ng/releases/download/0.9.0/raxml-ng_v0.9.0_linux_x86_64.zip):

```
raxml-ng --evaluate --msa alignment.phy --tree GTR20.treefile --model LG4X --threa
ds 4
```

### Likelihood mapping

This section benchmarks the likelihood mapping analysis in IQ-TREE 2 and TREE-PUZZLE.

For DNA analyses:

```
iqtree -s alignment.phy --prefix lmap_1 -m HKY -lmap 10000 -n 0 -T 1
iqtree -s alignment.phy --prefix lmap_4 -m HKY -lmap 10000 -n 0 -T 4
```

For Amino acid analyses:

```
iqtree -s alignment.phy --prefix lmap_1 -m LG -lmap 10000 -n 0 -T 1
iqtree -s alignment.phy --prefix lmap_4 -m LG -lmap 10000 -n 0 -T 4
```

We downloaded TREE-PUZZLE version 5.3.rc16 from [http://www.tree-puzzle.de](http://www.tree-puzzle.de). Note that TREE-PUZZLE only accepts alignment files in strict Phylip format (i.e., at most 10 characters in the sequence names).

1. Run `puzzle alignment.phy` to start a text menu.
2. Enter `b` to change `Type of analysis` to `Likelihood mapping`.
3. Enter `v` to change `Quartet evaluation criterion` to `Exact maximum likelihood`.
4. Enter `n` to change the `Number of quartets` to 10000.
5. Enter `y` to start the analysis.

### Fast tree search

This section benchmarks the `-fast` option in IQ-TREE 2 and FastTree 2.

For DNA analyses:

```
iqtree -s alignment.phy --prefix fast -m GTR+G -fast
iqtree -s alignment.phy --prefix normal -m GTR+G
```

For Amino Acid analyses:

```
iqtree -s alignment.phy --prefix fast -m LG+G -fast
iqtree -s alignment.phy --prefix normal -m LG+G
```

We downloaded FastTreeMP from [http://www.microbesonline.org/fasttree/](http://www.microbesonline.org/fasttree/).

For DNA analyses:

```
OMP_NUM_THREADS=1 /usr/bin/time FastTreeMP -nt -gtr -gamma alignment.fa >fasttree.
tree 2>fasttree.log
```

For Amino Acid analyses:

```
OMP_NUM_THREADS=1 /usr/bin/time FastTreeMP -nt -lg -gamma alignment.fa >fasttree.t
ree 2>fasttree.log
```

### Branch tests

This section benchmarks the SH-aLRT test in IQ-TREE 2 and PhyML.

For DNA analyses:

```
iqtree -s alignment.phy -T 4 --prefix alrt -m HKY+G -te GTR.treefile -alrt 1000
```

For Amino acid analyses:

```
iqtree -s alignment.phy -T 4 --prefix alrt -m LG+G -te GTR20.treefile -alrt 1000
```

We downloaded PhyML version 3.3.20190321 from [https://github.com/stephaneguindon/phyml/releases](https://github.com/stephaneguindon/phyml/releases) and followed the instructions to compile it.

For DNA analyses:

```
phyml-3.3 -i alignment.phy -b -4 -a e -u GTR.treefile -o r  --no_memory_check
```

For Amino Acid analyses:

```
phyml-3.3 -i alignment.phy -b -4 -a e -u GTR20.treefile -o r  --no_memory_check -d
 aa
```

### Tree topology tests

This section benchmarks the tree topology test implementation in IQ-TREE 2 and CONSEL. We first ran IQ-TREE 2 with `-wt` option to obtain a set of trees.

For DNA analyses:

```
iqtree -s alignment.phy -m JC -wt --prefix trees -T 10 -t PARS
head -n100 trees.treels >trees.100
```

For Amino Acid analyses:

```
iqtree -s alignment.phy -m LG -wt --prefix trees -T 10 -t PARS
head -n100 trees.treels >trees.100
```

We then performed all the tests in IQ-TREE 2 and also use `-wsl` option to write site log-likelihoods to `.sitelh` file for use in CONSEL later.

For DNA analyses:

```
iqtree -s alignment.phy -m GTR+G -te GTR.treefile -z trees.100 -zb 10000 -zw -au -
wsl -T 1 --prefix AU_1
iqtree -s alignment.phy -m GTR+G -te GTR.treefile -z trees.100 -zb 10000 -zw -au -
wsl -T 4 --prefix AU_4
```

For Amino Acid analyses:

```
iqtree -s alignment.phy -m LG+G -te GTR20.treefile -z trees.100 -zb 10000 -zw -au
-wsl -T 1 --prefix AU_1
iqtree -s alignment.phy -m LG+G -te GTR20.treefile -z trees.100 -zb 10000 -zw -au
-wsl -T 4 --prefix AU_4
```

We downloaded CONSEL version 0.20 from http://stat.sys.i.kyoto-u.ac.jp/prog/consel/, edited `src/Makefile` file to un-comment the line: `CFLAGS = -O3 -Wall`, so that the code will be compiled with optimization flag, and run `make` to compile CONSEL. This will generate several executables in the `src` folder, including `makermt`, `consel` and `catpv`. We run CONSEL with (the same for DNA and AA analyses):

```
/usr/bin/time makermt --puzzle AU >AU.makermt 2>&1
/usr/bin/time consel AU >AU.consel 2>&1
catpv -s 1 AU > AU.catpv
```
